# Immune-Coagulation Dynamics in Severe COVID-19: Insights from Autoantibody Profiling and Transcriptomics

**DOI:** 10.1101/2025.03.14.643398

**Authors:** Anoop T Ambikan, Axel Cederholm, Sefanit Rezene, Maribel Aranda-Guillen, Hampus Nordqvist, Carl Johan Treutiger, Ronaldo Lira-Junior, Nils Landegren, Soham Gupta

## Abstract

Severe COVID-19 is characterized by immune dysregulation and coagulation abnormalities, leading to complications such as thromboembolism and multi-organ failure. This study explores the relationship between autoantibodies targeting coagulation-related factors and gene expression in severe COVID-19. Whole-blood transcriptomics revealed upregulation of coagulation-related genes, including VWF and Factor V, in severe patients compared to mild cases and healthy controls. Autoantibody profiling against seven coagulation-related proteins (ADAMTS13, Factor V, Protein S, SERPINC1, Apo-H, PROC1, and Prothrombin) showed reactivities below established positivity thresholds, but mean-fluorescent intensities were elevated numerically in severe (Protein S) and convalescent (SERPINC1) patients. Correlation analysis revealed trends of negative associations between autoantibody reactivities and coagulation gene expression in severe cases, suggesting a potential role for autoantibodies in modulating immune-coagulation interactions warranting further orthogonal validation. Furthermore, age-dependent increases in subthreshold autoantibody reactivities were observed in severe cases, highlighting the potential impact of immunosenescence on disease severity. These findings do not exclude the possibility that subthreshold autoantibodies may contribute indirectly to immune-coagulation dynamics in severe COVID-19 through mechanisms beyond direct transcriptional regulation. This study highlights the complexity of immune-coagulation interactions and provides foundation for future research into their biological and clinical relevance, particularly for identifying biomarkers and therapeutic targets in thromboinflammatory diseases.

## INTRODUCTION

Coronavirus disease 2019 (COVID-19) is not only a respiratory illness but also a systemic condition in some instances characterized by significant immune dysregulation and hypercoagulability, particularly in severe cases. The severe manifestations of COVID-19 are occasionally driven by an excessive immune response coupled with a hypercoagulable state, both triggered by SARS-CoV-2 infection. This hypercoagulable state, characterized by coagulopathy and thrombosis, contributes to systemic microangiopathy, thromboembolism, and ultimately multi-organ failure [1, 2].

The interplay between inflammation and coagulation in COVID-19 is a key factor in disease severity. SARS-CoV-2 infection occasionally induces a robust inflammatory response in some patients, often described as a cytokine storm, which includes elevated levels of IL-6, TNF-α, and IL-1β. These cytokines activate the coagulation cascade, particularly the extrinsic pathway through tissue factor expression on endothelial cells and monocytes, leading to fibrin formation [3]. Simultaneously, the inflammatory response suppresses natural anticoagulant mechanisms, including antithrombin (SERPINC1) and the protein C system (Factor V and Protein S), further promoting a prothrombotic state [4].

This bidirectional relationship between inflammation and coagulation potentially creates a vicious cycle that exacerbates disease severity. For instance, thrombin (prothrombin), beyond its role in coagulation, also activates protease-activated receptors (PARs) on endothelial cells and platelets, further fueling inflammation [5]. The interplay between immune, complement and coagulation systems is a critical factor in some adverse outcomes of SARS-CoV-2 infection, contributing to both inflammation and thrombosis [6].

Emerging evidence suggests that autoantibodies, particularly those targeting type I interferons and other molecules, play a significant role in COVID-19 and are associated with adverse clinical outcomes [7-9]. Specifically, Zuo et al. (2020) reported that approximately 50% of serum samples from COVID-19 patients were positive for antiphospholipid autoantibodies [9]. Specific autoantibodies, such as those targeting prothrombin, have been detected following SARS-CoV-2 infection and are influenced by the strength of the antibody response against viral proteins, further implicating their role in COVID-19 severity [10]. Furthermore, injecting IgG fractions from these patients into mouse models resulted in enhanced venous thrombosis, highlighting the complex interplay between autoimmunity, inflammation, and coagulation in COVID-19 [9].

Given this intricate relationship, it is crucial to investigate how autoantibodies targeting coagulation-related factors may influence gene expression profiles, particularly in severe COVID-19. Autoantibodies such as those against ADAMTS13, Factor V, Protein S, SERPINC1, Apo-H, PROC1, and prothrombin could be hypothesized to exacerbate the dysregulation of the coagulation system, further intensifying the inflammatory response and leading to worse clinical outcomes. For example, autoantibodies against ADAMTS13, which regulates von Willebrand factor [11], can impair their normal functions, thereby increasing thrombotic risk as seen in COVID-19 patients who exhibit a higher prevalence of ADAMTS13 antibodies and markedly reduced ADAMTS13 activity compared to healthy individuals [12].

Despite growing evidence of the role of autoantibodies in COVID-19, a critical gap remains in understanding their correlation with disease severity and coagulation abnormalities. Transcriptomic analyses have identified distinct gene expression signatures linked to COVID-19 severity, and considering the presence of specific autoantibodies may help to elucidate this [13, 14]. To bridge these knowledge gaps, this study sets out to explore the relationship between autoantibodies targeting coagulation-related factors and COVID-19 severity by integrating autoantibody profiling with transcriptomic analysis. Our aim is to uncover novel insights into how gene expressions and autoantibodies influence COVID-19 pathogenesis, potentially leading to the identification of biomarkers for disease severity and the discovery of new therapeutic targets.

## RESULTS AND DISCUSSION

### Gene Expression in Coagulation and Complement Cascades in COVID-19 Patients

We reanalyzed the whole-blood transcriptomics data targeting the coagulation and complement cascade genes from our earlier study, which included 21 COVID-negative controls (HC), 10 convalescent, 28 patients with mild disease and 11 with severe disease [15]. Analysis was performed on 85 genes belonging to the ‘Complement and Coagulation Cascades’ in KEGG pathway (KEGG: 04610). The PCA plot shows varied expression across different health conditions, with notable differences between healthy controls, convalescent individuals, and those with mild or severe disease (**Figure 1A**). The heatmap shows that gene expression levels were elevated in COVID-19 patients, with a particularly pronounced increase observed in those with severe disease. As shown in **Figure 1B**, several genes in the coagulation cascade, such as VWF, PROS1 and Factor V, were significantly upregulated in severe disease as compared to mild disease state. However, heterogeneity in the gene expression pattern was observed, as not all patients demonstrated elevated levels, highlighting variability in the response among individuals.

**Figure 1:**
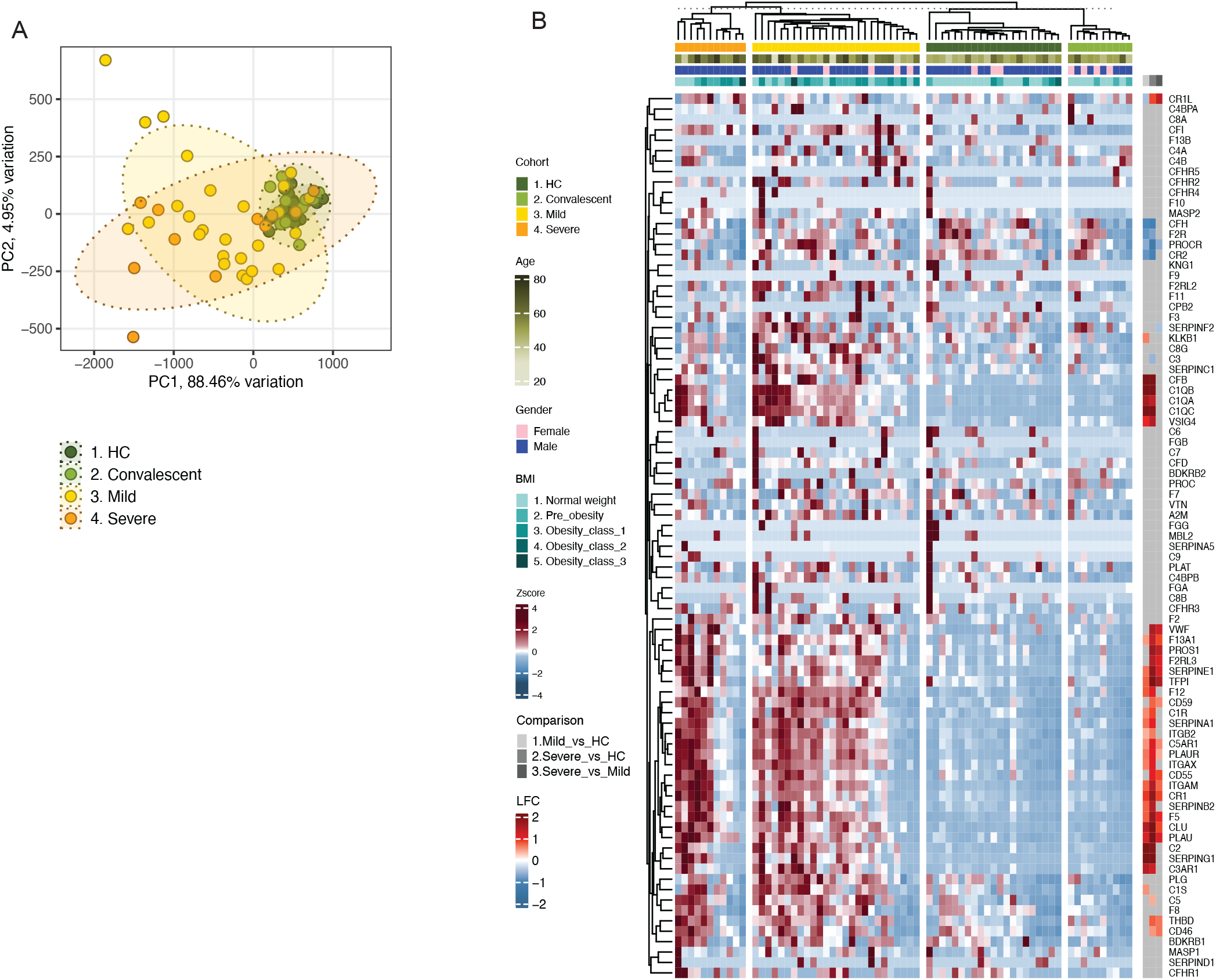
Gene Expression Analysis in COVID-19 and Control Groups. **A)** Principal Component Analysis (PCA) displaying the sample distribution based on the expression data of 85 genes within the ‘Complement and Coagulation Cascades’ pathways across 21 COVID-negative controls, 10 convalescent, 28 mild, and 11 severe COVID-19 cases. Samples are visually grouped by disease severity, illustrating distinct clusters, with PC1 accounting for 88.46% of variation and PC2 for 4.95%. **B)** Heatmap showing expression profile of genes involved in the coagulation and complement pathway across different patient groups. Color intensity of the heatmap encodes z-score normalized expression levels, accentuating marked upregulation in severe cases compared to others. Comparative analysis includes mild vs. healthy control (HC), severe vs. HC, and severe vs. mild, with expression fold changes annotated. Statistical significance was calculated using DESeq2 in R, with p-values adjusted for multiple comparisons using the Benjamini-Hochberg method.

The significant upregulation of several key genes involved in these pathways, such as VWF and Factor V, in patients with severe COVID-19 compared to those with mild disease underscores the central role of coagulation dysregulation in the pathogenesis of severe disease. This aligns with previous studies that have highlighted the hypercoagulable state as a hallmark of severe COVID-19, leading to thromboembolic events and multi-organ failure [1, 2]. However, the observed heterogeneity among individuals with severe disease suggests that additional factors, such as autoantibodies, may modulate the response to SARS-CoV-2 infection and contribute to disease variability.

### Autoantibody Profiling in COVID-19 Patients

The observed variability in gene expression patterns led us to investigate the presence of autoantibodies against coagulation factors, as they may contribute to the differential responses observed in COVID-19 patients. Using an in-house autoantibody-screening assay, we assessed the presence of circulating autoantibodies to seven coagulation-related factors (ADAMTS13, Factor V, Protein S, SERPINC1, Apo-H, PROC1, and prothrombin) and their relationship to the expression of genes associated with complement and coagulation pathway and to disease severity. **Figure 2** shows the mean fluorescent intensity (MFI) of each of the specific autoantibodies in different disease categories. The autoantibody reactivities detected for coagulation-related proteins were low compared to those normally observed for viral proteins and established autoantibody targets using this type of assay. Applying a positivity threshold defined either as the MFI of the healthy control (HC) group plus seven times their standard deviation (marked by the yellow dotted line in Figure 2) or an arbitrary cutoff of 1000 MFI (marked by blue dotted line in Figure 2) whichever was higher, our analysis did not reveal autoantibody positivity for any of the tested COVID-19 patients or controls. Therefore, we acknowledge that there is uncertainty about whether these subthreshold signals correspond to very low-level binding events or reflect non-specific background. Moreover, potential antigen-related assay limitations including cross-reactivity cannot be excluded. Further investigation, including more sensitive assays and orthogonal confirmatory methods, will be required to clarify if there is any biological relevance of these novel observations. This suggests that while autoantibodies against these coagulation factors are not universally elevated, the presence of such autoantibodies does not appear to be a common feature across all cases.

**Figure 2:**
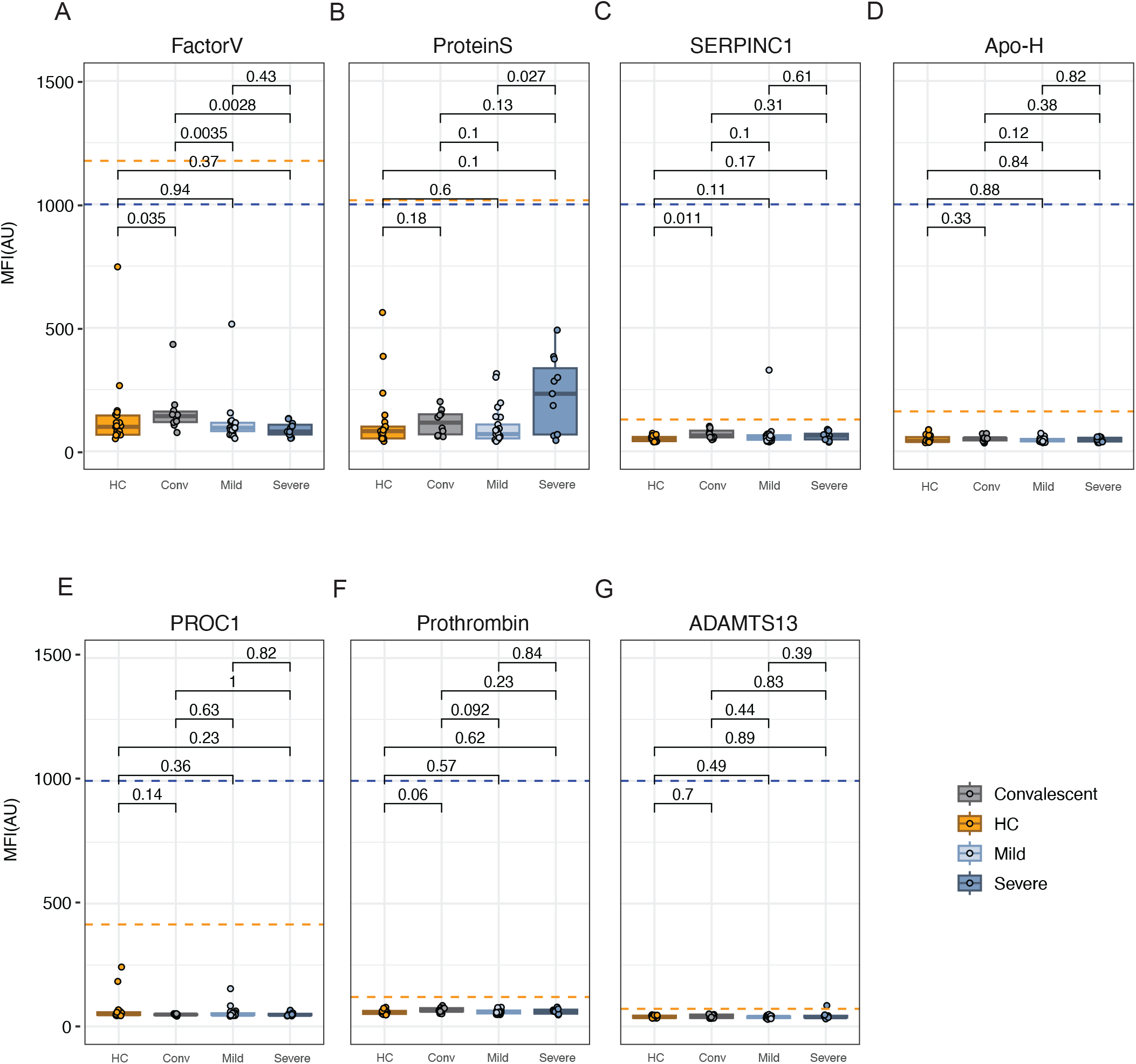
Coagulation related autoantibody Profiling in COVID-19 Patients. Analysis of autoantibodies against ADAMTS13 **(A)**, Factor V **(B)**, Protein S **(C)**, SERPINC1 **(D)**, Apo-H **(E)**, PROC1 **(F)**, and Prothrombin **(G)** depicted through dot plots for four groups: Healthy Controls, Convalescent, Mild, and Severe COVID-19 patients. Positivity thresholds are marked by yellow dotted lines, set at either the mean fluorescence intensity (MFI) plus seven standard deviations of the healthy control group (MFI+7SD) or an arbitrary cutoff of 1000 MFI denoted by blue dotted lines, whichever is higher. All samples are subthreshold for all antigens, including ADAMTS13, SERPINC1 and PROC1. Autoantibody reactivities were determined using a custom-developed Luminex assay. Statistical significance was evaluated using Mann-Whitney U tests, with p-values indicated for each comparison.

In pairwise comparisons of the subthreshold MFIs of each of the signals of autoantibody candidates across groups, convalescent patients exhibited significantly higher expression of anti-Factor V compared to both the active COVID-19 groups and healthy controls (HC) (Mann-Whitney U test, p<0.05). Patients with severe disease exhibited higher MFI levels of anti-protein S compared to the patients with mild disease (Mann-Whitney U test, p=0.027; **Figure 2B**) despite sub threshold reactivities. Interestingly, the subthreshold MFI signals of SERPINC1 and anti-prothrombin candidate autoantibodies were numerically elevated in convalescent individuals compared to healthy controls (p-values of 0.011 and 0.06, respectively; Figure 2C and 2F. This observation aligns with prior studies suggesting an enrichment of these autoantibodies following SARS-CoV-2 infection [10]. Differences in the applied statistical tests indicate the need for further investigations into autoantibodies in these groups, and these screening assay results does not contradict previous reports suggesting that autoantibodies can exacerbate the inflammatory and coagulation responses in COVID-19, potentially leading to more severe outcomes [7, 9]. These observations underscore the potential for further exploratory studies to investigate autoantibodies and prevalence post-infection, emphasizing the need for a detailed analytical approach across different patient groups.

### Correlation Between Autoantibody Data and Gene Expression

We subsequently analyzed correlations between the expression of coagulation-related genes and MFIs of the autoantibody data across different patient groups, despite no patient meeting the established positivity cut-off. This exploratory approach aimed to identify subtle patterns that, while not clinically significant, may offer insights into the underlying biological processes of COVID-19. Notably, within the healthy controls (HC) and convalescent groups, correlations were minimal (Figure 3A-B), which is consistent with the expected stability and regulatory effectiveness of their immune systems. Conversely, the severe COVID-19 cases showed a trend of more notable R values (Figure 3D), hinting at potential interactions between specific autoantibodies and gene expression that could exacerbate disease severity through engagement with multiple biological pathways. However, it is important to mention that these correlations did not reach statistical significance after adjusting the p-values and should be interpreted cautiously (Supplementary Table 1). In contrast to severe cases, the mild COVID-19 group exhibited only limited correlations between gene expression and autoantibody reactivities (Figure 3C). The disparate correlation patterns observed between mild and severe COVID-19 cases leads us to speculate unique underlying pathogenic mechanisms tailored to the intensity of the disease. In severe cases, the predominantly negative correlations might reflect a tipping point where homeostatic mechanisms are overwhelmed, leading to uncontrolled coagulation and systemic inflammation. This scenario indicates a possible over-activation of immune responses that could drive the severity of clinical manifestations. Conversely, the milder cases display a pattern of more favorable or neutral correlations, hinting at a more regulated and balanced interplay between the immune and coagulation systems. This controlled response potentially curtails the escalation of the disease, helping to maintain physiological stability and prevent the progression to severe forms of the illness. The lack of significant adjusted p-values, despite noticeable R values, suggests that while trends in the data exist, they do not reach statistical significance. This highlights the need for cautious interpretation of these correlations and further investigation into their biological impact on disease progression in COVID-19.

**Figure 3:**
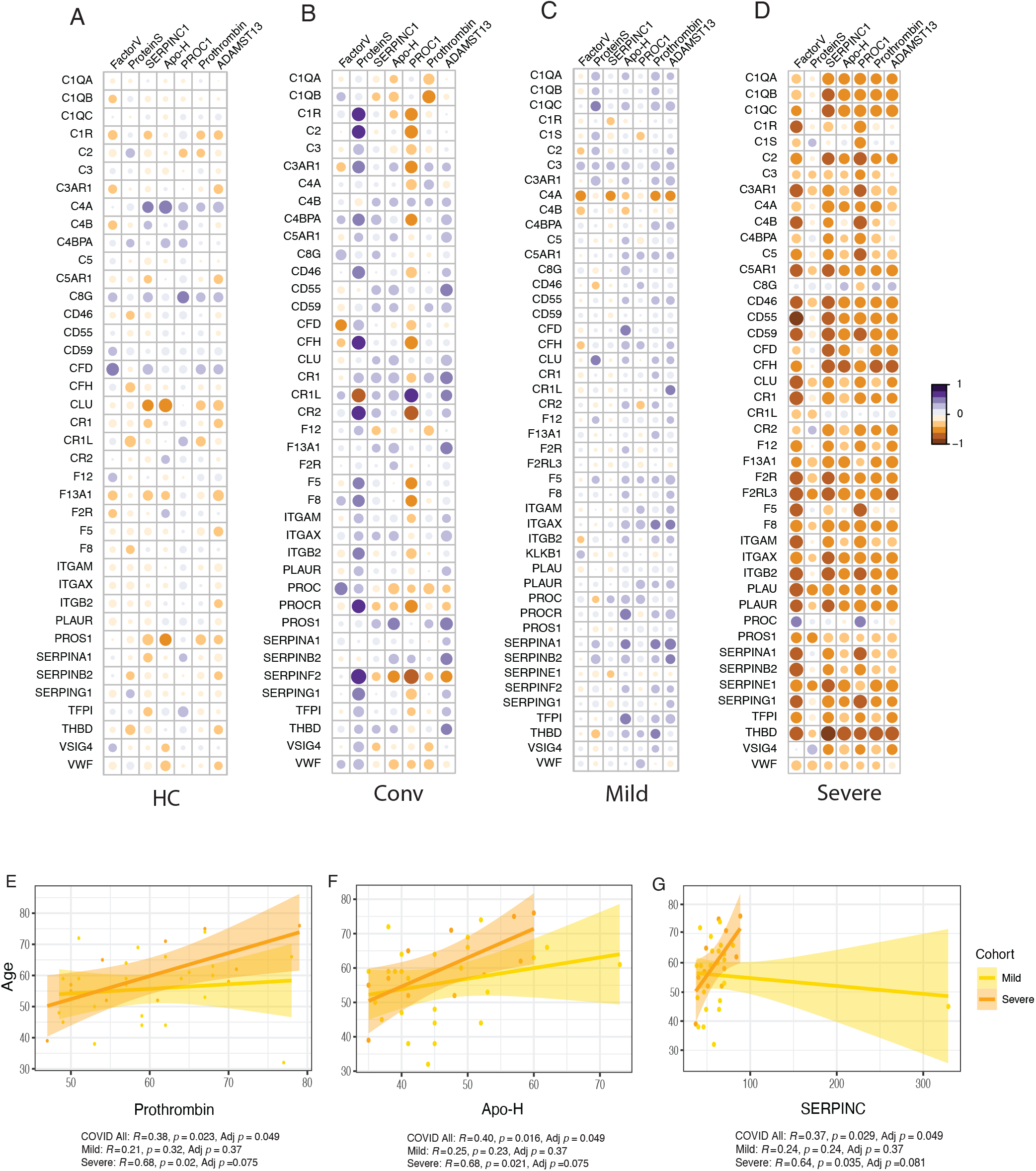
Correlation Analysis Between Autoantibody data and Gene Expression. **A-D)** This figure presents correlation heatmaps (Corr-plot) for Healthy Controls **(A)**, Convalescent **(B)**, Mild COVID-19 cases **(C)**, and Severe COVID-19 cases **(D).** Each bubble illustrates the relationship between subthreshold autoantibody data and the expression of genes within the coagulation and complement pathways. The Spearman correlation coefficients are depicted in each cell, with color intensity reflecting the strength and direction of correlations (red for positive, blue for negative). Analyses were conducted using the psych package in R. **E-G)** Scatter plots depicting the correlation between age and subthreshold MFI of autoantibodies targeting Prothrombin **(A**), Apo-H **(B**), and SERPINC1 **(C)**, across all COVID-19 patients and separated by severity (Mild: Yellow and Severe: Orange). Each point represents an individual patient; lines indicate the best fit trend between the two features with shaded areas for the 95% confidence interval. Spearman’s correlation coefficients, p-values and adjusted p-values are provided, with adjustments for multiple testing using the Benjamini-Hochberg method.

Interestingly, although the subthreshold MFI levels of Protein S autoantibodies were higher in severe cases than in mild cases (Figure 2B), the correlations with gene expression changes were poor (Figure 3D). This lack of correlation suggests that the pathogenic role of these autoantibodies, if any, may be independent of direct transcriptional changes in coagulation-related genes, potentially acting through post-translational modifications or interactions with other immune pathways that were not captured in this analysis. This complexity underscores that autoantibodies might influence disease severity through multiple, possibly non-genomic, mechanisms, indicating the possibility of a sophisticated interplay of immune responses in COVID-19 pathogenesis.

Gene expression of Thrombomodulin (THBD1), a key regulator of thrombin homeostasis [16] showed a pronounced negative correlation with SERPINC1 (R = -0.88), Prothrombin (R = - 0.779), Factor V (R = -0.745), ADAMTS13 (R = -0.763), and Apo-H (R = -0.776) autoantibody candidates in the severe group irrespective of their MFI levels being below the positivity threshold, with all correlations exceeding an R value of -0.75 and p< 0.01, showing a clear trend, although the adjusted p-values did not fall below the conventional threshold for significance (Adj-p< 0.25) (Figure 3D). These patterns suggest a potential disruption in the normal regulatory mechanisms of coagulation and anticoagulation under severe disease conditions. Conversely, in milder cases, THBD1 exhibited a positive correlation trend (R = 0.41) with prothrombin autoantibodies (**Figure 3C**), potentially indicating a more controlled or compensatory immune response that may help maintain some degree of homeostasis. These contrasting correlation patterns between mild and severe COVID-19 cases could suggest potential differences in the underlying pathogenic mechanisms driving disease severity influenced by THBD1’s regulation of coagulation dynamics across different severities of COVID-19 and underscores the need of future investigations.

### Age-Dependent Autoantibody Reactivity

Our study further explored the relationship between age and autoantibody responses across different severities of COVID-19 for the sub threshold signals for autoantibody reactivities. In severe cases, we observed strong positive correlations between age and the subthreshold MFI of autoantibodies against Prothrombin (R = 0.68, p = 0.02, adj-p = 0.075), Apo-H (R = 0.68, p = 0.021, adj-p = 0.075), and SERPINC1 (R = 0.64, p = 0.035, adj-p = 0.081) (**Figure 3E-3G**), suggesting an increased immune response in older individuals. These correlations were more pronounced compared to mild cases, where the relationships were weaker and not statistically significant after adjustment. Interestingly, when analyzing all COVID-19 patients together, despite lower correlation coefficients, certain subthreshold autoantibody candidates such as ADAMTS13 (R = 0.375, p = 0.024, adj-p = 0.049) and Apo-H (R = 0.397, p = 0.016, adj-p = 0.049) exhibited statistically significant adjusted p-values (Supplementary Table 2). This indicates that while individual correlations of sub threshold reactivities may appear modest, collectively they reveal significant age-related trends across the COVID-19 spectrum. Although, these findings emphasize the complex influence of age on immune system behavior, suggesting that older patients might have distinct immune profiles that could impact their response to COVID-19, it is important to note that the MFI levels did not surpass our detection threshold, hence these results should be interpreted with caution. This observation aligns with studies suggesting that aging is associated with an increase in autoantibody production due to immunosenescence and inflammaging, contributing to the exacerbated clinical manifestations of diseases like COVID-19 [8]. For example, older COVID-19 patients frequently exhibit a more severe disease course, characterized by elevated autoantibody levels that may enhance hypercoagulability and increase the risk of thromboembolic events [9]. The correlation of these autoantibodies with disease severity underscores the need for age-specific therapeutic strategies that address both immune dysregulation and the heightened coagulation state observed in the elderly [17]. The higher prevalence of autoantibodies in older patients [18] accentuates the need to integrate immune modulation and anticoagulation therapies into COVID-19 management strategies specifically designed for this demographic. Future research should delve deeper into how aging affects autoantibody production including new-onset ones that emerges following infection [19] and its implications for disease progression and therapeutic outcomes in elderly COVID-19 patients.

Our research contributes to the growing body of knowledge on the critical role of autoantibodies in the pathophysiology of COVID-19, especially in severe cases. We observed that while there are correlations between autoantibody data and coagulation gene expression alterations, the detected autoantibody reactivities were below the positivity thresholds, possibly suggesting potential roles in disease pathology that warrants further investigation. This finding highlights the complexity of immune interactions in COVID-19 and supports the need for further studies to elucidate the mechanisms by which autoantibodies affect disease progression and outcomes, especially in long-term sequelae showing persistent complement dysregulation and thromboinflammation, as recently noted by Cervia-Hasler et al. (2024) [20]. Understanding these interactions could lead to the development of targeted therapies and biomarkers for disease severity, enhancing our management strategies for both acute and prolonged manifestations of COVID-19.

In conclusion, this study underscores the potential of incorporating both autoantibody profiling and gene expression analysis to dissect the complex pathogenesis of severe COVID-19. While the direct impact of autoantibodies on disease processes requires further validation, our findings hint at a potential role in modulating disease severity, suggesting that they might influence a range of pathophysiological pathways. These insights may inform future studies and therapeutic strategies aimed at modulating immune responses to improve clinical outcomes in severe COVID-19 cases. Further research will be crucial to establish the clinical relevance of these autoantibodies and their utility in guiding interventions for COVID-19, particularly in severe and long-term scenarios.

## METHODOLOGY

### Study Cohort

The cohort of COVID-19 patients and healthy controls used in this study has been previously described in detail [21]. Briefly, the study included 41 hospitalized COVID-19 patients who tested positive for SARS-CoV-2, stratified by oxygen consumption into two groups: mild (O2 consumption <4 l/min; n=28) and severe (O2 consumption ≥4 l/min; n=11).

Exclusion criteria included significant pre-existing conditions (liver cirrhosis, severe renal insufficiency, chronic obstructive pulmonary disease, and chronic lung diseases leading to habitual SpO2 ≤ 92%). Additionally, 31 healthy controls (HC) were included, with 10 testing positive for SARS-CoV-2 antibodies, referred to as Convalescent (Conv). All study participants provided informed consent, with procedures approved by the regional ethics committees of Stockholm (dnr 2020-01865) and conducted in accordance with the Declaration of Helsinki.

### Autoantibody Detection

The methodology for detecting human IgG autoantibodies, previously described [22], includes the use of magnetic beads (MagPlex®, Luminex Corp.) prepared using the AnteoTech Activation Kit for Multiplex Microspheres to ensure effective coupling with specific antigenic proteins. The beads were coupled with commercial proteins, including ADAMTS13, Factor V, Protein S, SERPINC1, Apo-H, PROC1 and Prothrombin, at standardized coupling ratios (1.5 × 10^6 beads per 3 μg of protein). Stored plasma samples obtained from the patients were initially diluted in PBS (1:25), followed by a secondary dilution (1:10) in PBS with 0.05% Tween, 3% BSA, and 5% Milk. The bead-sample mixture was then incubated at 650 rpm for two hours. Following three wash cycles in PBS with 0.05% Tween, the beads were fixed in 0.2% PFA for 10 minutes. Detection was carried out using a secondary antibody from Invitrogen, with incubation lasting 30 minutes before analysis on a FlexMap 3D instrument (Luminex Corp).

### Transcriptomics Data Analysis

Whole blood transcriptomics data for the analysis was sourced from the previously published study by the group [15]. The data were reanalyzed, focusing on differential gene expression related to the coagulation and complement cascades. Genes associated with these pathways, part of the KEGG pathway gene set (KEGG: 04610), were retrieved from the Enrichr libraries [23]. The transcriptomics data for the analysis were sourced from this previously published study. Differential gene expression analysis was conducted using the R package DESeq2 v1.44.0 [24]. To adjust for potential biases, characteristics such as age, gender, BMI, and other unwanted variations, as identified by the RUVSeq package v1.38.0 [25], were included in the model matrix. Genes with an adjusted p-value (p_adj) less than 0.05 and a log2 fold change greater than one was considered significantly regulated. Principal component analysis (PCA) was performed with the PCAtools package v2.16.0 in R, using transcripts per million (TPM) normalized data from the complement and coagulation cascade genes. The Wilcoxon test was executed using the stat_compare_means function in R. Spearman’s correlation was calculated using the corr.test function from the psych package v2.4.3. Genes with a variance across samples below 0.1 were excluded from the correlation analysis. Corrections for multiple hypothesis testing were applied using the Benjamini-Hochberg method following the correlation analysis. Visualization of PCA dimensions and the generation of dot plots were carried out using the ggplot2 package v3.5.1 in R. The heatmap was generated using the ComplexHeatmap package v2.20.0, and the corrplot was created with the corrplot package v0.92.

## FIGURE LEGENDS

**Supplementary Figure 1:** Extended Analysis of Subthreshold Autoantibody Distributions Comprehensive heat map representation of subthreshold autoantibody reactivities, stratified by various patient demographics (age, gender, and BMI) and clinical categories. Subthreshold reactivities for each antigen is analyzed across different patient groups, with color codes and intensities indicating the relative reactivities.

**Supplementary Table 1:** Correlation between subthreshold autoantibody reactivities (MFI) and complement and coagulation gene set normalized expression values (TPM). Genes with variance less than 0.1 were omitted.

**Supplementary Table 2:** Correlation between subthreshold autoantibody reactivities (MFI) and age in all COVID patients.

## Acknowledgements

The authors would like to thank the excellent support received from the study nurses Elisabet Storgard and Ronnie Ask, Södersjukhuset. We thank the National Facility for Autoimmunity and Serology Profiling at SciLifeLab for excellent technical support related to the autoantibody experiments. Additionally, we are grateful to Dr. Ujjwal Neogi, Karolinska Institutet for setting up the cohort and sharing the data, which was invaluable for this study. This work was funded by the Swedish Research Council grant to S.G. (2021-03035) and N.L. (2021-03118), and the Center for Medical Innovation (CIMED-FoUI093304), along with additional support from the Karolinska Institutet Stiftelser och Fonder (grant 2022-02232) to S.G. S.R. received support through the KID funding program (S.G). N.L. acknowledges support from the Göran Gustafsson Foundation (2141 and 2227), while R.L.R. received support from SOF/Stockholm County Council (grant FoUI-966258).

## Author Contributions

S.G. conceptualized and with help from N.L supervised the study. A.T.A. performed the bioinformatics analysis, data interpretation and figure generation. A.C. performed the autoantibody profiling and contributed to data analysis. S.R. and M.A.-G. contributed to experimental work and data interpretation. H.N. and C.J.T. managed patient cohort data and provided clinical insights. R.L.J. contributed to the experimental design and critical review of the manuscript. N.L. assisted with autoantibody assay development and provided expertise in immunological profiling. S.G., R.L.J, A.T.A and A.C wrote the manuscript, with critical input from all authors. All authors have read and approved the final version of the manuscript.

## Data Limitations and Perspectives

This study provides insights into the interplay between coagulation gene expression and autoantibody data in severe COVID-19. However, the absence of significant autoantibody positivity limits the ability to draw definitive conclusions. The findings suggest a potential role in modulating disease severity based on subthreshold autoantibody reactivities and gene expressions data, which requires validation in larger cohorts. Future studies should explore post-translational modifications and broader immune-coagulation dynamics to unravel the complex mechanisms underlying severe COVID-19.

## Data Availability Statement

The datasets generated and analyzed during the current study are available from the corresponding author on reasonable request.

## Ethics Approval Statement

The study was approved by the Regional Ethics Committees of Stockholm (Dnr 2020-01865) and conducted in accordance with the Declaration of Helsinki.

## Patient Consent Statement

Informed consent was obtained from all participants.

