## Supplementary figures and images for "Immune-Coagulation Dynamics in Severe COVID-19: Insights from Autoantibody Profiling and Transcriptomics"

### Supplemental Figure

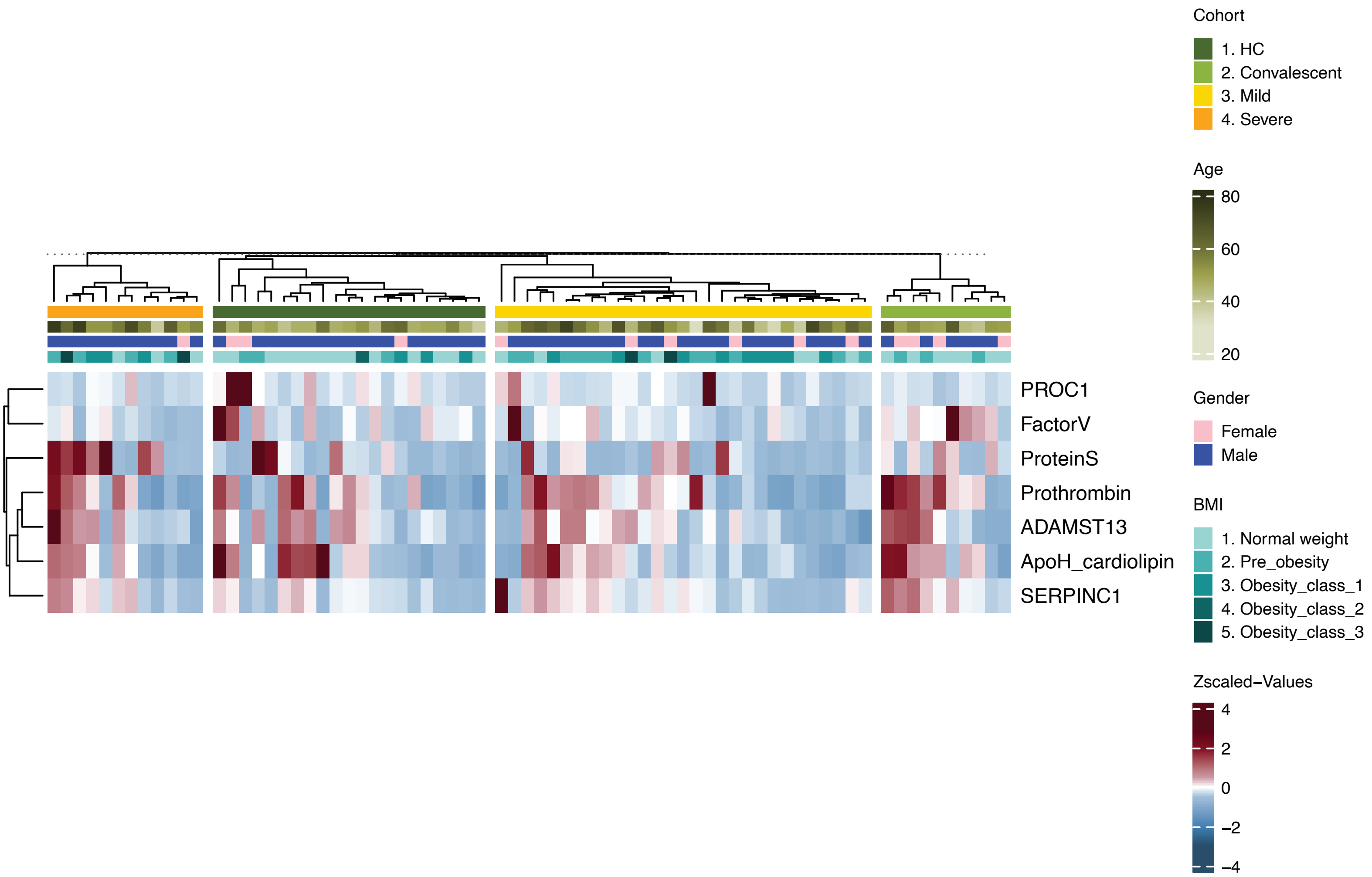
